# FtsZ phosphorylation brings about growth arrest upon DNA damage in *Deinococcus radiodurans*

**DOI:** 10.1101/2022.06.16.496254

**Authors:** Reema Chaudhary, Shruti Mishra, Ganesh Kumar Maurya, Hari S Misra

## Abstract

The polymerization/depolymerization dynamics of FtsZ plays the pivotal role in cell division in majority bacteria. *Deinococcus radiodurans*, a radiation resistant bacterium, shows an arrest of growth in response to DNA damage, despite no change in the level of FtsZ. This bacterium does not deploy LexA/RecA type of DNA damage response and cell cycle regulation, and its genome does not encode homologs of *E. coli*’s SulA, which attenuate FtsZ functions in response to DNA damage in different bacteria. A radiation responsive Ser/Thr quinoprotein kinase (RqkA) characterized for its role in radiation resistance in this bacterium, could phosphorylate several cognate proteins including FtsZ (drFtsZ) at Serine 235 (S235) and Serine 335 (S335) residues. Here, we report the detailed characterization of S235 and S335 phosphorylation effect in the regulation of drFtsZ functions, and demonstrated that the phospho-mimetic replacements of these residues in drFtsZ had grossly affected its functions that could result in cell cycle arrest in response to DNA damage in *D. radiodurans*. Interestingly, the phospho-ablative replacements were found to be nearly similar to drFtsZ while phospho-mimetic mutant showed the loss of signatures characteristics to wild type enzyme including the arrest in its dynamics under normal conditions. The post-bleaching recovery kinetics for drFtsZ and phospho-mimetic mutant was nearly similar at 2h post irradiation recovery but found to be different under normal conditions. These results highlighted the role of S/T phosphorylation in the regulation of drFtsZ functions and cell cycle arrest in response to DNA damage and is first time demonstrated in this prokaryote.

## Introduction

The bacterial cell division is orchestrated by the coordinated functions of an array of cell division proteins called the divisome and genome segregation proteins called the segrosome. The mechanistic details of the processes of cell division have been extensively studied in both Gram-negative and Gram-positive bacteria like *Escherichia coli, Bacillus subtilis, Streptococcus pneumoniae, Staphylococcus aureus, Mycobacterium tuberculosis* and *Caulobacter crescentus* (Adams & Errington, 2009). The formation of divisome at the mid-cell position is both spatially and temporally regulated (Haeusser et al., 2016; Du & Lutkenhaus, 2019). FtsZ, a tubulin homolog and an important protein in the bacterial cell division, undergoes GTP-dependent polymerization-depolymerization dynamics, and it forms FtsZ-ring (Z-ring) largely at mid cell position (de Boer et al., 1992; Bramhill and Thomson, 1994; Oliva et al., 2004). FtsZ exists in two conformations: an open conformation observed in the filaments and a closed conformation in the monomeric form (Wagstaff et al., 2017). It is found to be a key protein that plays a crucial role in bacterial cell division. The properties of FtsZ protein are influenced by other proteins through protein-protein interactions. For example, FtsA, ZipA and Zap proteins help in anchoring and stabilizing the Z-ring at the mid-cell position, the MinCDE, SlmA and Noc proteins prevent the formation of Z-ring at the undesired sites *albeit* through different mechanisms. The stability of Z-ring at the cytoplasmic membrane requires interaction with other proteins, and also acts as a scaffold for the recruitment of downstream proteins for septal peptidoglycan biosynthesis (Lutkenhaus, 2007; Erickson et al., 2017). The GTP binding pocket is found to be conserved in FtsZ across different bacterial species and is required for the polymerization of FtsZ. The dynamics of FtsZ polymerization /depolymerization play a crucial role in the bacterial cell division (Bi & Lutkenhaus, 1993; Lutkenhaus, 2007; Dajkovic et al., 2008; Chen et al., 2012; Bisson-Filho et al., 2015). In response to DNA damage, the functions of FtsZ are attenuated by SOS response proteins like SulA in *E. coli* (Mukherjee et al., 1998), YneA in *B. subtilis* (Kawai et al., 2003) and Rv2719c in *M. tuberculosis* (Chauhan et al., 2006).

*Deinococcus radiodurans*, an extraordinary radiation-resistant bacterium that divides in perpendicular planes. It can tolerate the mutagenic effects of several DNA damaging agents including radiation and desiccation (Battista, 2000; Cox and Battista, 2005). The *D. radiodurans* shows growth arrest in response to γ irradiation (Modi et al., 2014). The transcriptome analyses showed the changes in the expression kinetics of DNA repair genes (Liu et al., 2003; Tanaka et al, 2004) and protein recycling in cells recovering from γ irradiation (Basu and Apte, 2012). Notably, there was no change in the levels of some putative cell division proteins including FtsZ during post irradiation recovery (PIR) (Modi et al., 2014). *D. radiodurans* lacks the LexA/RecA type DNA damage response and cell cycle regulation (Narumi et al., 2001) and its genome does not encode FtsZ attenuator like SulA or its homologs. How this bacterium regulates FtsZ function and cell cycle is intriguing. A γ radiation responsive Ser/Thr quinoprotein kinase encoded on an ORF DR_2518 (named as RqkA) has been characterized in this bacterium and its roles in radiation resistance and DNA double-strand break (DSB) repair have been demonstrated (Rajpurohit and Misra, 2010). RqkA phosphorylates a large number of cognate proteins including those involved in DNA repair and cell division, and thus a possibility of RqkA regulating the global response of DNA damage and cell cycle would be interesting to investigate. Here, we provided evidence that the phosphorylation of deinococcal FtsZ (drFtsZ) at serine 235 (S235) and 335 (S335) positions during PIR and by RqkA attenuates the functions of drFtsZ including the arrest of its polymerization/depolymerization dynamics in *D. radiodurans*. Although, FtsZ phosphorylation has been reported in canonical SOS response conferring bacteria, the role of S/T phosphorylation in FtsZ by a DNA damage responsive STPK and its effect in the γ radiation stressed growth is first time shown in any bacteria. The findings reported in this work may form a basis for proposing an STPK based mechanism of DNA damage response and the cell cycle regulation in prokaryotes also.

## Materials and methods

### Bacterial strains, plasmids and materials

*Deinococcus radiodurans* R1 (ATCC13939) was a kind gift from Professor J. Ortner, Germany (Schaefer et al., 2000). *Escherichia coli* strains DH5α and NovaBlue were used for cloning and maintenance of the plasmids, and *E. coli* strain BL21 (DE3) pLysS was used for the expression of recombinant proteins. *E. coli* was grown in Luria Bertini (LB) broth and *D. radiodurans* was grown in TGY medium (1% Bacto tryptone, 0.1% glucose, 0.5% yeast extract) with shaking at 180 rpm at 37 ºC and 32 ºC, respectively. *E. coli* harboring the recombinant expression vectors and their derivatives were grown in the presence of antibiotics, as required. Molecular biology grade chemicals and enzymes were purchased from Merk Inc. and New England Biolabs, USA.

### Site directed mutagenesis using overlap-PCR

Phospho-site mapping of drFtsZ exposed to RqkA kinase has shown that serine residues located at 235 (S235) and 335 (S335) positions are phosphorylated (Maurya et al., 2018). For studying the effect of phosphorylation on the functional aspects of drFtsZ, the S235 and S335 were substituted with alanine (phospho-ablative) or aspartic acid (phospho-mimetic) in different combinations (Table 1). Overlapping primers were designed to mutate S235 of drFtsZ. The *ftsZ* gene segments were amplified using two sets of primers; forward *ftsZ* and reverse *ftsZ*^S235A^, and forward *ftsZ*^S235A^ and reverse *ftsZ* primers in separate reactions. These gene segments were used as a template for another PCR to generate a full-length *ftsZ* gene carrying the desired mutation using the *ftsZ* forward and reverse primers including the restriction sites. For hybridization and an extension of the overlapping strands, two PCR cycles were run as follows: denaturation (3 min), 20 cycles of denaturation (30 sec, 95 ºC), annealing (2 min, required temperature), extension (1 min 10 sec, 72 ºC) followed by an extension of 5 min at 72 ºC and then reaction was subjected to another PCR cycle of denaturation (2 min), 20 cycles of denaturation (30 sec, 95 ºC), annealing (35 sec, required temperature), extension (1 min 10 sec, 72 ºC) followed by an extension of 10 min at 72 ºC. The *ftsZ* mutant alleles were cloned in pET28a (+) at *Nde*I and *Bam*HI yielding pETZS235A, pETZS253D, pETZS335A and pETZS335D. The double phospho-mutants were generated using pETZS335A and pETZS335D as templates using overlapping primers as shown in Table 1 and double mutants bearing recombinant plasmids like pETZAA, pETZAD, pETZDA and pETZDD were obtained (Table 1). The recombinant plasmids were further transformed into *E. coli* BL21 cells and the protein expression was ascertained by SDS-PAGE.

**Table 1:**
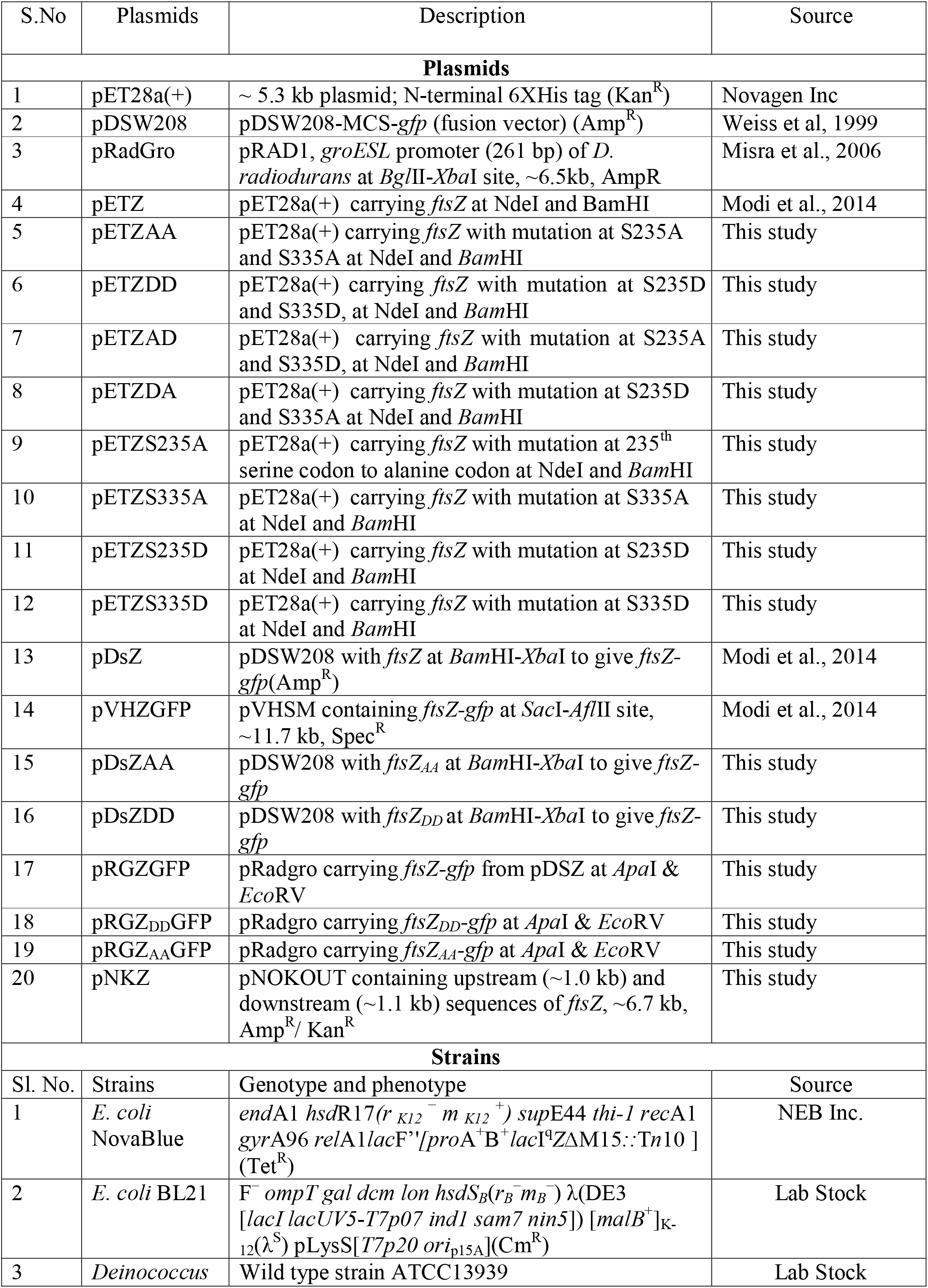
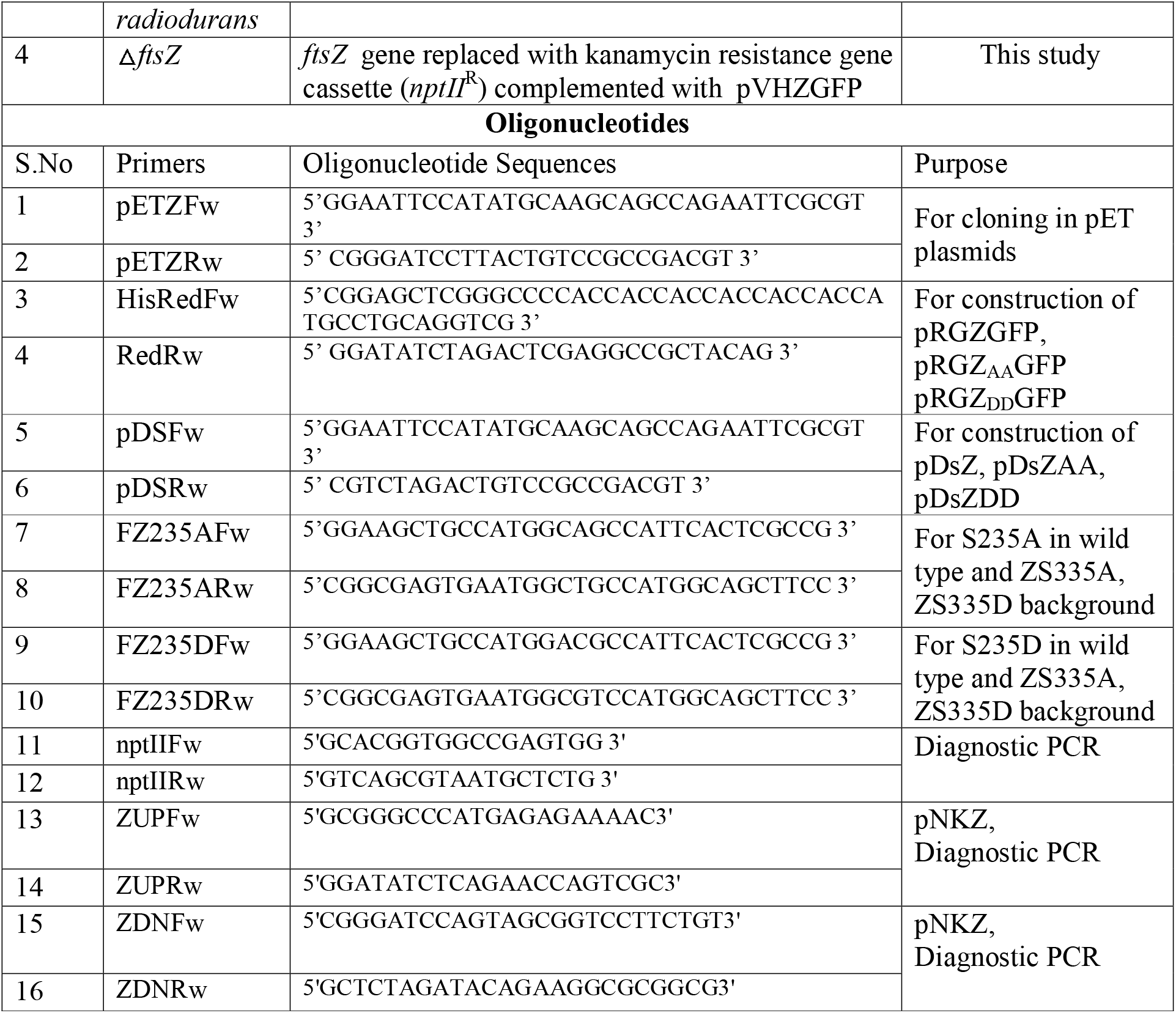
List of plasmids, strains and primers used in this study.

The coding sequences for drFtsZ, FtsZ_AA_ and FtsZ_DD_ were PCR amplified using the required primers from pETZ, pETZAA and pETZDD, respectively. These products were cloned in pDSW208 (Weiss et al., 1999) at *Bam*HI and *Xba*I yielding pDsZ, pDsZAA and pDsZDD. The coding sequence for translation fusion of drFtsZ and FtsZ_DD_ with GFP were PCR amplified from pDsZ and pDsZDD, respectively. These PCR products were cloned in pRadgro (Misra et al., 2006) at *Apa*I-*Eco*RV yielding pRGZGP and pRGZDDGFP. These recombinant plasmids are further transformed in *D. radiodurans* using the protocol as mentioned in Chaudhary et al., 2021.

### Purification of proteins

The recombinant drFtsZ (∼39kDa) and its mutant derivatives expressing in the *E*.*coli* BL21 cells were purified to near homogeneity using protocol as described (Modi et al., 2014). In brief, an overnight grown culture expressing the recombinant protein was diluted at 1:100 in fresh LB broth containing 25 μg/ml kanamycin. To this, 0.5 mM isopropyl-β-D-thiogalactopyranoside (IPTG) was added in exponentially growing cells and these were harvested 3 h post-induction. Cell pellet was suspended in buffer A (20 mM Tris–HCl, pH 7.6, 250 mM NaCl) containing 10 mM imidazole, 0.5 mg/ml lysozyme, 1 mM PMSF, 0.03 % NP-40, 0.03 % Triton X-100 and 10 % glycerol) and incubated at 37 ºC for 30 min. To this, the protease inhibitor cocktail was added and the cells were sonicated for 10 min at 15s pulses with intermittent cooling for 5 min at 25 % amplitude. The cell lysate was centrifuged at 11,000 rpm for 30 min at 4 ºC. The cell-free extract was loaded onto NiCl_2_ charged fast-flow chelating sepharose column (GE Healthcare) pre-equilibrated with buffer A. The column was washed with 20 column volumes of buffer A containing 50 mM imidazole till proteins stop coming from the column. Recombinant proteins were eluted with buffer A containing 100, 200, 250- and 300 mM imidazole. The fractions were analyzed by SDS-PAGE and 250 mM and 300 mM fractions containing nearly pure proteins were pooled and concentrated using 10 kDa cut-off spin columns and then centrifuged at 14,000 rpm for 60 min to remove aggregates. The concentrated protein is dialysed against buffer A which was further subjected to purification using the gel-filtration sepharose column (GE healthcare) and collected using an automated AKTA collector. The fractions were analyzed by SDS-PAGE and those containing nearly pure proteins were pooled and histidine tag was removed with Factor Xa using manufacturer’s protocol. The mixture was concentrated using 10 kDa cut-off spin columns and then centrifuged at 14,000 rpm for 60 min. The protein in supernatant was dialyzed in buffer A containing 10 mM Tris-HCl pH 7.6, 250 mM NaCl, 20% glycerol and 1 mM PMSF and stored at −20°C.

### Biochemical assays

The GTPase activity was measured as the release of [^32^P]-αGDPs from [^32^P]-αGTPs by thin-layer chromatography (TLC). In brief, proteins in different concentrations were pre-incubated in 20 µl polymerization buffer (PB) (20 mM Tris-HCl, pH 7.5, 75 mM KCl) supplemented with MgCl_2_ as required at 37 ºC for 10 min before the reaction was initiated with [α^32^P]-GTP. Reaction mixtures were further incubated at 37°C for 20 min. 2 µl of the reaction mixture was spotted on the PEI Cellulose F+TLC sheets. Spots were dried and mixtures were separated on the solid support in a buffer (0.75 M KH_2_PO_4_/H_3_PO_4_ (pH 3.5)). TLC sheets were exposed to X-ray films in a light-proof cassette. Autoradiograms were developed and images were analyzed using Image-J and Graphpad Prism 6 software. The GTPase activity was also measured using the malachite-green assay as per the manufacturer’s protocol (Sigma-MAK113). The binding of GTP to recombinant proteins was checked as described in (Rohn, 1999). In brief, 100 nM 2′/3′-O-(N-methyl-anthraniloyl) guanosine 5′ triphosphate, trisodium salt (Mant-GTP) (M12415; Invitrogen) was incubated with 5 µM of wild type and mutant proteins separately in binding buffer (Tris-HCl, pH 7.6; 50 mM, KCl; 30 mM and MgCl_2_; 0.5 mM) and Mant-GTP binding to protein was monitored as the emission using JASCO spectrofluorometer FP-8500.

### Biophysical characterization

The quality of purified recombinant proteins was assayed by Circular Dichroism (CD) spectroscopy as described earlier (Maurya et al., 2019). In brief, 0.5 µM protein was diluted in a buffer containing 20 mM Tris (pH 7.6) and 150 mM NaCl and the spectra were recorded using a CD spectrophotometer (Biologic spectrometer MOS-500).

Further, the sedimentation assay was carried out using a modified protocol as described earlier (Modi et al., 2014). In brief, 5 µM of the purified recombinant drFtsZ and phospho-mutant derivatives were pre-incubated in polymerization buffer (PB) supplemented with 5 mM MgCl_2_ followed by the addition of 1 mM GTP/GDP as required. In all these experiments, the protein was pre-incubated in PB buffer with 5 mM MgCl_2_ at 37 ºC for 5 min and 1mM GTP or GDP and further incubated for 20 min at 37 ºC. The reaction mixtures were centrifuged at 14,000 rpm for 30 min at room temperature. The pellet and supernatants were separated and analyzed on 12% SDS-PAGE. Protein bands were visualized by Coomassie brilliant blue staining. Images were analyzed using Image-J and Graphpad software.

The polymerization dynamics of phospho-mutants of drFtsZ were monitored by 90° dynamic light scattering (DLS) as described in (Charaka and Misra, 2012). In brief, all the solutions used in this study were passed through a 0.22 µm filter and protein was centrifuged at 14,000 rpm for 60 min at 4 ºC. The DLS spectra were recorded with 5 µM purified proteins incubated with and without 1 mM GTP in PB buffer for 5 min at 37 ºC using Malvern Zetasizer. Data was recorded for 20 min at an interval of 30 sec and light scattering intensity in kilocounts per sec (kcps) was plotted against time using Graphpad software.

For transmission electron microscopy, 5 µM of wild type and mutant proteins were incubated in polymerization buffer (PB buffer; 50 mM Tris-HCl, pH 7.6, 50mM KCl and 5 mM MgCl_2_) for 5 min at room temperature followed by 15 min incubation at 37 °C in the presence of 1 mM GTP. For mounting, the carbon-coated TEM grids were charged under UV and reaction mixtures were put on the grids followed by the stain (w/v: 2% uranyl acetate). The grids were washed with Milli-Q water and left for drying at room temperature overnight and observed at a magnification of 10,000x (120 keV) using a transmission electron microscope JEM-1400 (JEOL).

### Confocal microscopy

Confocal microscopy was performed on an IX3SVR motorized platform using an Olympus IX83 inverted microscope with the laser beams focused on the back focal plane of a 100 × 1.40 NA oil-immersion apochromatic objective lens (Olympus, Inc). The intensity and time sequence of laser illumination at the sample was tuned using an installed Fluoview^**TM**^ software. The series of Z-planes were acquired at every 400-nm. Fluorescence emission was collected through a DM-405/488 dichroic mirror and the corresponding single-band emission filters.

For fixed-cell imaging, the bacterial cells were grown to mid-log or late-log phase in LB or TYG broth, fixed with 4 % paraformaldehyde for 10 min on ice and washed two times with phosphate-buffered saline (PBS; pH 7.4). These cells were stained with DAPI (0.5 µg/µl) for 10 min on ice and then washed three times with phosphate buffer saline (PBS). After washing, the cells were re-suspended in PBS, mounted on a 1 % agarose bed on glass slides and samples were observed. For image analysis, the 200-300 cells were taken from at least two separate microscopic fields captured in two independent experiments and analyzed for required attributes. The image analysis and other cell parameters were measured using automated the cellSens software. Data obtained were subjected to Student *t*-test analysis using statistical programs of GraphPad Prism. For time-lapse microscopy, the cells in the exponential phase were rinsed in PBS before imaging. Cells were deposited on an agarose pad made with 2X-TYG and impregnated with air holes to oxygenate the cells. The 3-D images (Z-planes were acquired every 400 nm) were acquired every 45-60 min for a period of 4-5 h using very low 488 nm laser power. Images were processed using cellSens software and Adobe Photoshop 7.0.

For fluorescence recovery after photobleaching (FRAP)-experiments, deinococcal cells expressing pRGZGFP and pRGZDDGFP in exponential phase were exposed to 6 kGy dose of γ radiation using GC-5000 whereas control cells were kept on ice. After γ-irradiation, both un-irradiated (UI) and irradiated (I) cells expressing FtsZ_WT_ and FtsZ_DD_ were collected at 0h and 2h of post-irradiation recovery. The cells were processed for confocal microscopy as described above. The region of interest was exposed to a laser bleach pulse of 35 to 45 ms followed by the acquisition of recovery for 30-40 s at low laser power. The fluorescence intensities in the bleached and other regions at each time point were extracted using the cellSens software and exported to Excel 2013. For each spot, the background intensity was subtracted and a correction factor was applied for overall photobleaching of the region of interest during observation using easy-FRAP software. Recovery half-times were determined by performing a non-linear regression fit of the intensity of the bleached region over time using GraphPad software (Anderson et al., 2004).

### Immunoblotting of total cellular proteins

Immunoblotting of total proteins was carried out as described earlier (Chaudhary et al., 2021). In brief, *E. coli* cells expressing pDSW208, pDsZ, pDsZAA and pDsZDD were grown at 37°C. The optical density of bacterial cultures were adjusted to 1.0 and cells were centrifuged. The cell pellets were washed with 1X PBS and cells were lysed at 95 °C for 15 min in equal volume of 10 mM Tris-HCl pH8.0, 1 mM EDTA and 2X Laemmli buffer. The equal amount of protein was run on 12 % SDS-PAGE and blotted on PVDF membrane using BioRad semi-dry blot machine. Blots were hybridised with monoclonal antibodies against GFP followed by anti-mouse alkaline-phosphatase conjugated antibody and signals were developed using NBT-BCIP (Roche).

### Construction of conditional null mutant of ftsZ

The recombinant plasmid pNKZ was constructed for the deletion of *ftsZ* (DR_0631) from chromosome I in *D. radiodurans*, respectively. For that, ∼ 1 kb of upstream and downstream regions from the coding sequence of FtsZ was PCR amplified using sequence specific primers (Table 2). PCR products were digested with required restriction enzymes and cloned in the pNOKOUT plasmid. The upstream and downstream fragments of *ftsZ* were cloned at *Apa*I-*EcoR*I and *Bam*HI-*Xba*I sites in pNOKOUT (Kan^R^) to give pNKZ. For the expression of FtsZ *in trans*, the recombinant plasmid pVHZGFP expressing FtsZ-GFP under *“Spac”* promoter was used and replacement of chromosomal copy of *ftsZ* with *nptII* using pNKZ was achieved as described earlier (Chaudhary et al., 2021). The clones having replacement of chromosal copy of ftsZ in the presence of episomal copy of wild type protein was named as conditional null mutant of *ftsZ* (Δ*ftsZ*). The recombinant plasmids; pRGZGFP, pRGZ_AA_GFP and pRGZ_DD_GFP were also transformed into Δ*ftsZ* to monitor the effect of phosphor-ablative and phosphor-mimetic replacements on the physiological functions of FtsZ *in-vivo*. The wild-type and other derivatives expressing FtsZ on plasmids were grown in the presence of respective antibiotics and their growth at OD_600_ was monitored in sterile 24-well microtitre plate using Synergy H1 Hybrid multi-mode microplate reader, Biotek for overnight at 32 °C. The data was processed, analysed for the statistical significance and plotted using GraphPad Prism software.

## Results

### The serine residues at 235^th^ and 335^th^ are mutated to alanine and aspartate codons

The FtsZ homologs are largely conserved across the bacteria and are structurally divided into 3 domains; N-terminal domain, globular conserved core domain and C-terminal domain-containing linker and variable regions (Fig 1A (i)). Multiple sequence alignment of FtsZ from different bacteria showed that deinococcal FtsZ (drFtsZ) is ∼41.77% identical and ∼56.61% similar with FtsZ of *Bacillus subtilis* (Fig 1A (ii)). The positional conservation of serine residues at the 235^th^ and 335^th^ position of drFtsZ analysed across the FtsZ of different bacteria was found to be non-conserved (Fig 1A (iii)). The structure of drFtsZ was modelled using I-TASSER and refined using ModLoop and RAMPAGE server (Fig 1B (i)). The modelled structure was aligned with the structure of FtsZ of *B. subtilis* (2VXY) using TM-Align and Pymol (Fig 1B (ii)). The aligned models show an RMS value of 1.36 and a TM score of 0.96034 indicating that the two structures are highly identical. Earlier, the RqkA phosphorylation of S235 and S335 residues in drFtsZ has been demonstrated (Maurya et al., 2018). The placement of these residues in the 3-D structure of drFtsZ showed that the S235 and S335 residues are in close proximity to GTP binding pocket and in the intrinsically disordered peptide (IDP) region of the 3-D structure of drFtsZ, respectively. The S235 and S335 were replaced with alanine

**Fig 1:**
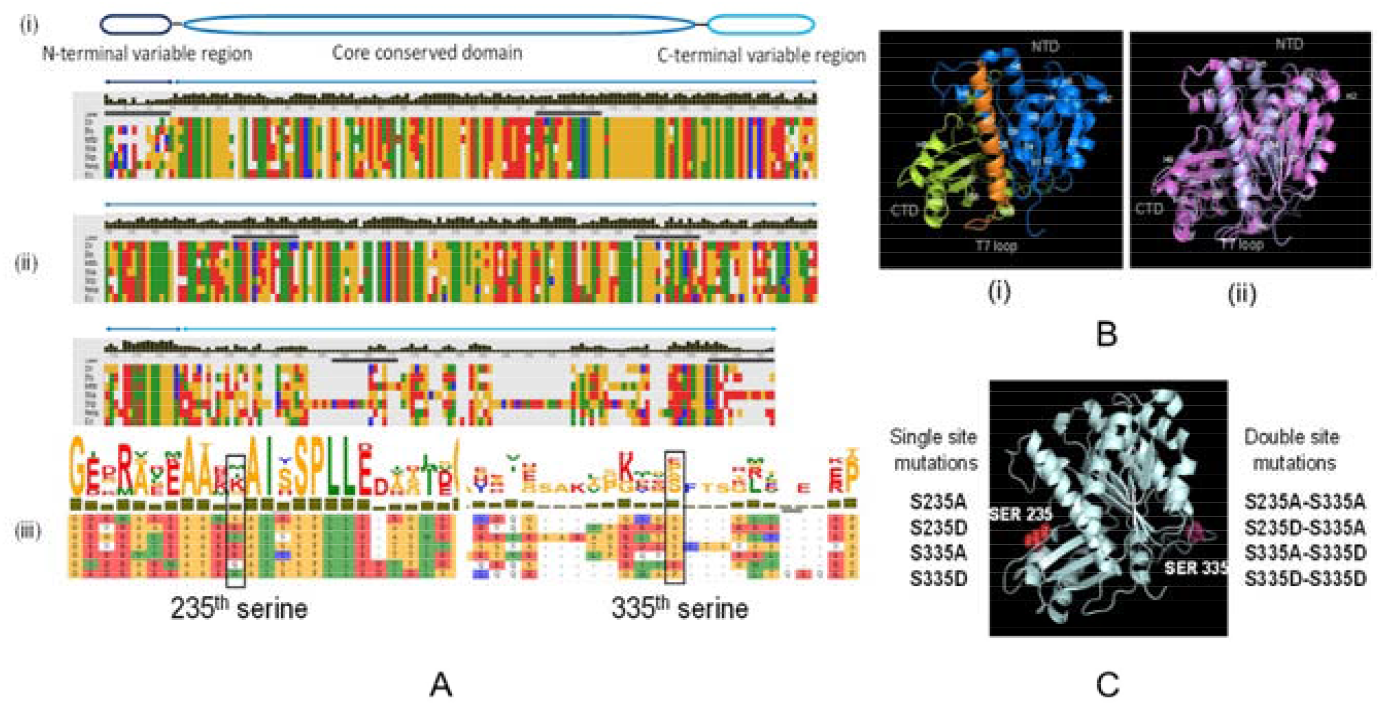
The sequence analysis, structural prediction and representation of mutations in drFtsZ. The general representation of the domains and regions of FtsZ (i), the multiple sequence alignment using sequence of FtsZ from different bacterial species is performed using MSAProbs webserver (ii), 235^th^ and 335^th^ serine residue are shown separately where it indicates less order of conservation at these residues across several bacterial species (iii) **(A)**. Structure of deinococcal FtsZ (drFtsZ) was modelled using I–Tasser and ModLoop server where NTD (N-terminal domain), T7-loop and CTD (C-terminal domain) are labelled (i), The structure of drFtsZ overlapped with bsFtsZ (*Bacillus subtilis* FtsZ; PDB ID 2VAM) (ii) **(B)**. The two serine which were mapped as phospho-sites in drFtsZ are mutated to alanine and aspartate codons in single and double combinations as shown in the figure **(C)**.

### Phospho-mimetic replacement of S235 and S335 affects GTPase activity and interaction with GTP

The recombinant drFtsZ and its mutants’ proteins were purified (Fig S1A) and checked for secondary structure alteration if any, using Circular Dichroism (CD) spectroscopy. All the mutant proteins produced the typical CD spectra nearly similar to drFtsZ with the clear dips at 208 and 222 nm suggesting that all have α-helix secondary structures typical to wild type protein (Figure S1B). The GTPase activity of S235 and S335 mutants were estimated using thin-layer chromatography and compared with drFtsZ (Fig 2A). Since the polymerization and GTPase activity of FtsZ requires a critical concentration of the protein, the activity was monitored at different protein concentrations *in vitro* for all the mutant forms. The replacement of S235 and S335 residues with aspartate has compromised the GTPase activity *albeit* to different levels. For instance, the FtsZ_DD_ mutant showed a significant loss in GTPase activity (∼ 3.0 ± 0.29-fold) as compared to wild type as well as the double phospho-ablative, S235AS335A (hereafter referred to as FtsZ_AA_) (Fig 2B). Furthermore, the other single phospho-mimetic/ablative and double mutants, having one substitution with phospho-ablative residue while the other with phospho-mimetic residue has also showed reduced GTPase activity. For instance, the single mutants like FtsZ_S235A_, FtsZ_S335A_, FtsZ_S235D_ and FtsZ_S335D_ showed respective GTPase activity by ∼1.67 ± 0.32, ∼1.62 ± 0.44, ∼4.27 ± 0.16 and ∼5.1 ± 0.188 fold less. However, the fold reduction in GTPase activity in double mutants FtsZ_AD_ and FtsZ_DA_ was ∼3.35 ± 0.58, ∼3.46 ± 0.38 fold, respectively as compared to drFtsZ at 5 µM. Interestingly, similar trends were obtained at 10 µM of protein concentration (data not shown). The loss of GTPase activity in mutants could be explained if these mutations have affected the wild type interaction of GTP and /or polymerization ability of these proteins.

**Fig 2:**
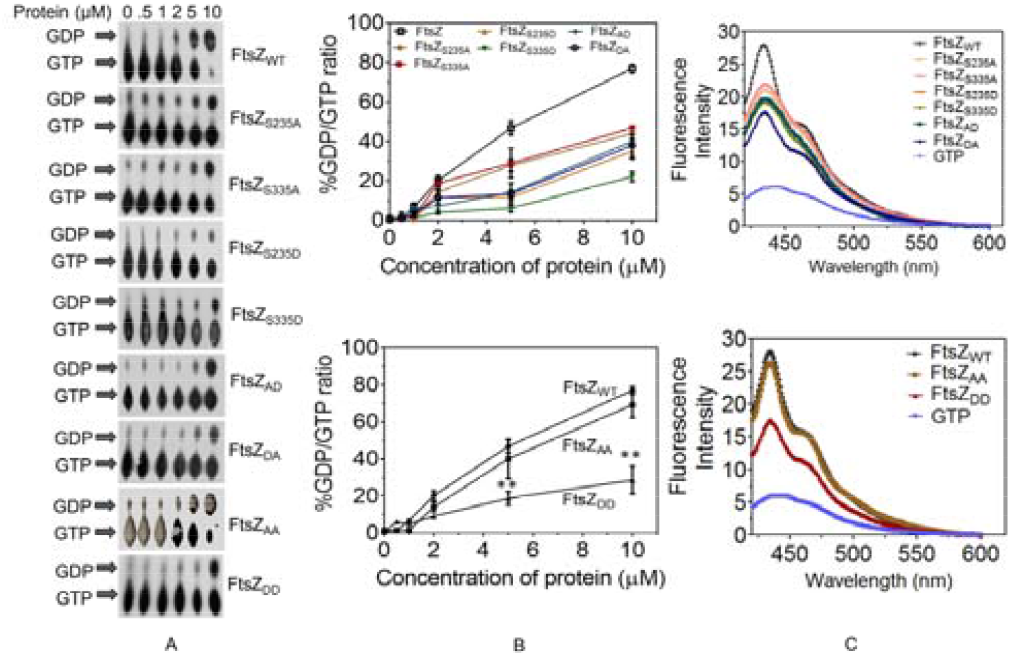
Characterization of phospho-mutants of drFtsZ for their GTPase activity and GTP-binding. Purified recombinant FtsZ_WT_ and its phospho-mutants as labelled in the figure were pre-incubated in polymerization (PB) buffer in a concentration dependent manner. Reaction mixtures were spotted on PEI Cellulose F+ TLC sheets, dried and separated in buffer system. TLC sheets were exposed to X-ray films and autoradiograms were developed and the conversion of GTP to GDP is marked with the arrows **(A)**. Images were analysed using ImageJ. The percentage of GDP/GTP was calculated for each concentration of the different proteins and plotted using GraphPad Prism **(B)**. Nucleotide binding assay of wild-type and mutants of FtsZ was also performed. Emission spectra for mant-GTP alone and bound to wild-type and different mutants of FtsZ were recorded from 410 to 600 nm at an excitation wavelength of 355 nm. The color coded lines represent the fluorescence signal from 100 nM mant-GTP in the presence of different proteins as mentioned in the figure **(C)**.

GTP binding assay was performed using 2′/3′-O-(N-methyl-anthraniloyl) guanosine 5′ triphosphate, trisodium salt (Mant-GTP) and 5 µM of wild type and other mutant proteins as described in methods. The increase in fluorescence signals of Mant-GTP was observed in the presence of all the proteins. However, FtsZ_S235A_, FtsZ_S335A_, FtsZ_S235D_, FtsZ_S335D,_ FtsZ_AD_, FtsZ_DA_ and FtsZ_DD_ mutants showed a relatively low increase in Mant-GTP fluorescence intensity as compared to wild type and FtsZ_AA_ (Fig 2C). This effect was more pronounced in FtsZ_DD_ mutant and surprisingly there was no change when both the residues were replaced with alanine in FtsZ_AA._ These results raised a strong possibility of S235 and S335 phosphorylation affecting drFtsZ affinity to GTP suggesting that the phosphorylation of these residues seems to have affected the catalytic GTPase activity by affecting the nucleotide interaction with mutant forms of drFtsZ.

### Phospho-mimetic replacement of S235 and S335 affects the polymerization ability of drFtsZ

Since the phospho-mimetic replacement of S235 and S335 has affected the GTPase activity and GTP interaction. Therefore, the effect of S235 and S335 phosphorylation on the polymerization of drFtsZ was examined using 90º angle dynamic light scattering (DLS), sedimentation assay and transmission electron microscopy. DLS analysis showed that the polymerization of the FtsZ_WT_ and all the phosphor-ablative mutants including FtsZ_AA_ was stimulated by the addition of GTP while there was not change in the size of polymers in phopsho-mimetic mutant including FtsZ_DD_ (Fig 3A). The light scattering signal remained near baseline for phosphor-mimetic mutants as compared to wild type and phosphor-ablative mutants. Estimation of polymer size through sedimentation at higher speed showed similar results as that of DLS. Results showed that the amount of the phosphor-mimetic mutants including FtsZ_DD_ in the pellet was found to be nearly ∼13.57 ± 2.48 times lesser than the drFtsZ in the presence of GTP and Mg^+2^. This indicated that the presence of GTP and Mg^+2^ could not induce the formation of higher order structures in phosphor-memetic mutants, which was observed in drFtsZ and phosphor-ablative mutants including FtsZ_AA_ (Fig 3B). Transmission electron microscopic studies also showed that the drFtsZ and phosphor-ablative mutants including FtsZ_AA_ can form higher order structures while phospho-mimetic derivatives including FtsZ_DD_ did not produce higher order structures (Fig 3C).

**Fig 3:**
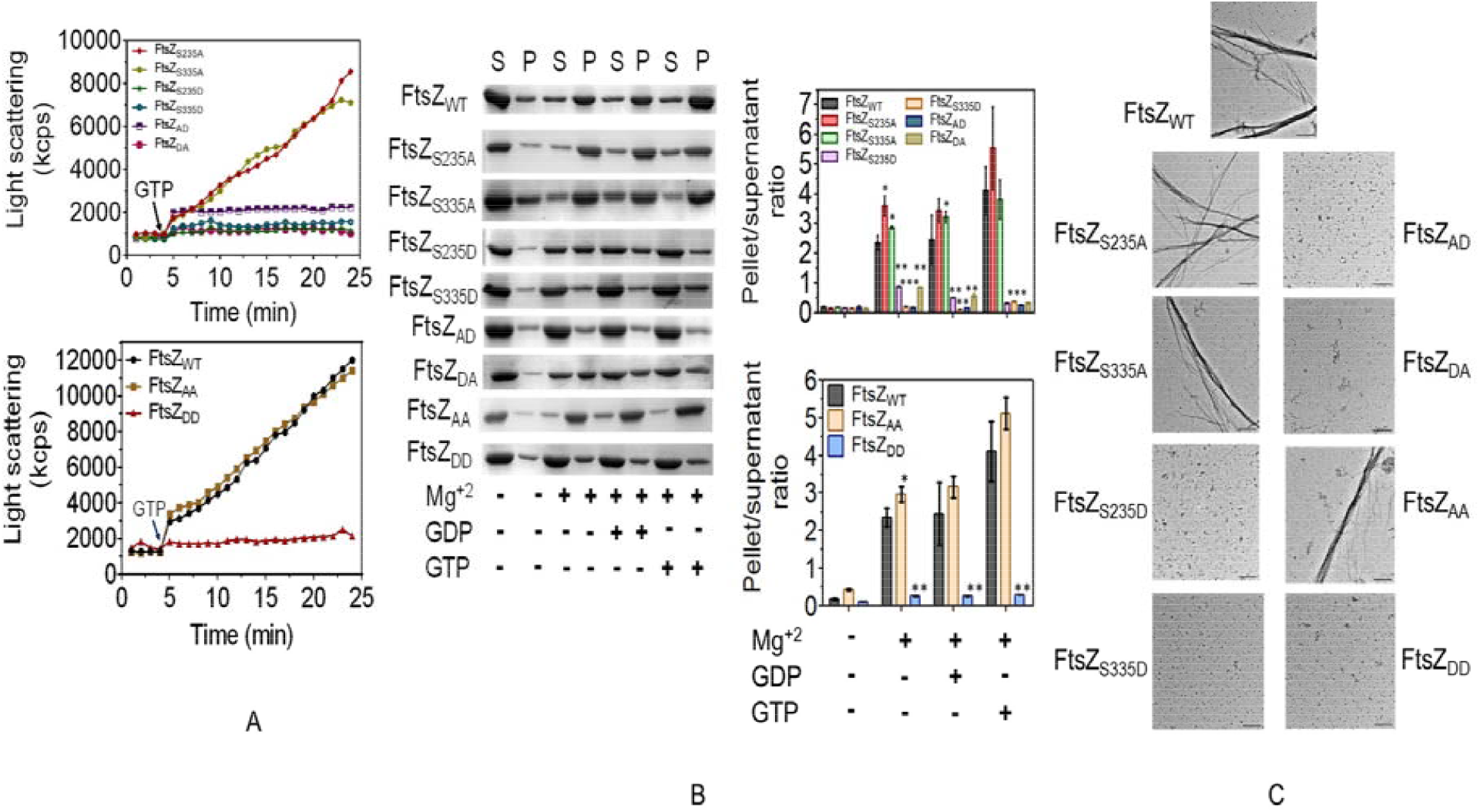
Monitoring of phospho-mutants of FtsZ for their polymerization abilities. 5µM of the purified recombinant FtsZ_WT_ and other phospho-mutant proteins were preincubated in PB buffer only followed by the addition of GTP (shown by arrow) at 37 ºC for 5 min in a quartz cuvette and then the light scattering was recorded for 20 min at an interval of 30 seconds. Light scattering intensity in kilocounts per sec (kcps) was plotted against time using GraphPad software for each protein as mentioned in the graph **(A)**. Similarly, 5µM of the purified recombinant FtsZ and phospho-mutant derivatives were preincubated in PB buffer supplemented with 5 mM MgCl_2_, followed by addition of 1mM of GTP/GDP and incubated for 20 minutes at 37 ºC. Reaction mixtures were centrifuged and the pellet (P) and supernatant (S) fractions were separated and analyzed on SDS-PAGE, stained with Coomassie brilliant-blue. Images were analyzed using Image-J. The ratio of pellet to supernatant was plotted using GraphPad software **(B)**. For transmission electron microscopy, 5 µM of wild-type and mutant proteins were incubated in polymerization buffer (PB buffer; Tris-HCl, pH 7.6; 50mM, KCl; 50mM and MgCl_2_; 1mM) for 5 min at room temperature followed by the addition of 1 mM GTP and incubation at 37 °C followed by mounting on carbon-coated TEM grids. The grids were observed using transmission electron microscope JEM-1400 (JEOL) transmission electron microscope with a magnification of 10,000x (Scale bar 500 nm) **(C)**.

### Phospho-mimetic replacement of S235 and S335 altered the physiological behaviour of drFtsZ

The expression of translation fusion of drFtsZ (pDsZ; Modi et al., 2014), FtsZ_AA_ (pDsZAA) and FtsZ_DD_ (pDsZDD) with GFP was ascertained in the *E. coli* cells using confocal microscope. The FtsZ_AA_ mutant protein formed the spiral-like cytoskeletal structures in the cells mocking the phenotype of cells over-expressing drFtsZ (Fig 4A). The *E. coli* cells expressing the drFtsZ and FtsZ_AA_ have shown the very long cells. A similar observation has been reported in the *E. coli* cells overexpressing its native FtsZ episomally (Dai and Lutkenhaus, 1992). Interestingly unlike the wild-type and FtsZ_AA_, the FtsZ_DD_ expressing *E. coli* cells have shown normal foci in normal size cells (Fig 4A). This phenotype had been attributed to the levels of protein in the recombinant *E. coli* cells (Dai and Lutkenhaus, 1992). Therefore, the levels of drFtsZ, FtsZ_AA_ and FtsZ_DD_ proteins were measured in the *E. coli* cells expressing these proteins from the same vector under similar promoter. All the types of cells like drFtsZ, FtsZ_AA_ and FtsZ_DD_ showed nearly same levels of proteins detected by drFtsZ antibodies (Fig S2), suggesting that mimicking phosphorylation in drFtsZ (FtsZ_DD_) alters its classical behaviour at least in *E. coli*.

**Fig 4:**
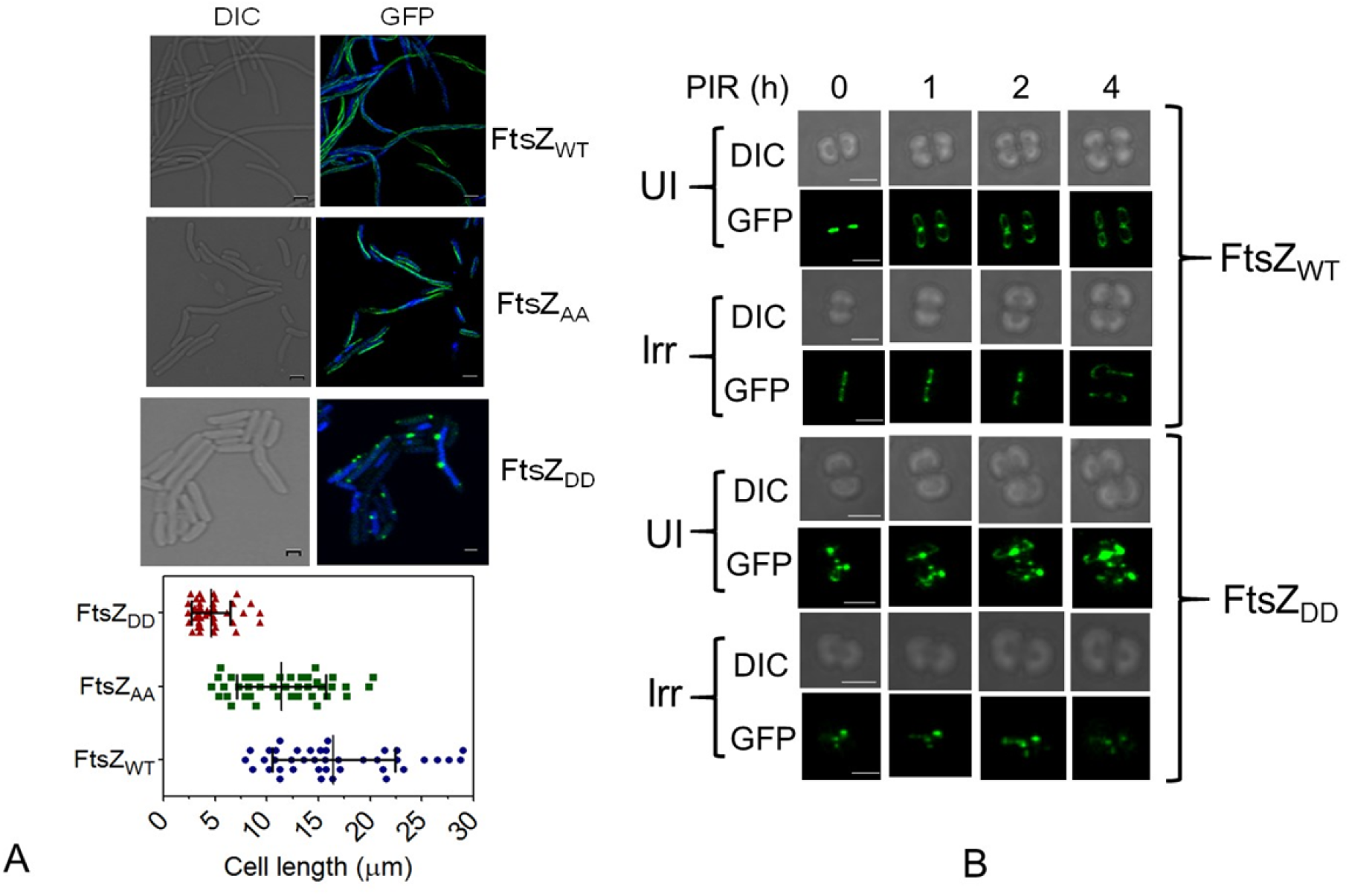
Effect of the mutations on drFtsZ localization dynamics and *in-vivo* functions. For fixed-cell imaging, the *E. coli* cells expressing pDSZ (FtsZ_WT_), pDsZAA (FtsZ_AA_) and pDsZDD (FtsZ_DD_) were fixed and observed using confocal microscope Olympus IX83 inverted microscope equipped with an IX3SVR motorized platform. These cells were also stained with DAPI. The image analysis was done using automated cellSens software. Data obtained were subjected to Student *t*-test analysis using statistical programs of GraphPad Prism. Scale bar denotes 2 µm **(A)**. For time-lapse imaging of the cell growth and division process, cells in exponential phase were exposed to 6 kGy dose of γ radiation whereas control cells were kept on ice. After γ-irradiation, both un-irradiated (UI) and irradiated (Irr) cells expressing pRGZGFP (FtsZ_WT_) (A) and pRGZ_DD_GFP (FtsZ_DD_) mounted on 2X-TYG agarose pad before imaging using confocal microscope Olympus IX83 inverted microscope. Air holes were designed in agarose pads to oxygenate the cells. The 3-D images (Z-planes were acquired every 400 nm) were acquired every 45-60 min for a period of 4-5 h using very low 488nm laser power. Images were processed using the cellSens software and Adobe Photoshop 7.0.

Subsequently, the localization pattern and dynamics of both drFtsZ (pRGZGFP) and FtsZ_DD_ (pRGZDDGFP) were monitored in *D. radiodurans* exposed to γ radiation. The cells expressing drFtsZ and FtsZ_DD_ were observed under time-lapse microscopy. drFtsZ localization and cellular dynamics were found to be different under normal and γ stressed conditions. For instance, under normal conditions, the drFtsZ-ring formation was found to be perpendicular to the previous plane of cell division in time scale that corresponds to 1-2 h PIR (Fig 4B) and the constriction of the cell envelope was evident by the closure and separation of Z-rings in the daughter cells in time scale that corresponds to 4h PIR (Fig 4B). Upon γ-radiation exposure, the formation of Z-ring in the required plane was observed in nearly 4h and onward PIR. Since, drFtsZ undergoes phosphorylation during this time scale of PIR, a possibility of delayed Z ring formation and dynamics due to phosphorylation of drFtsZ could be speculated. To answer it, the cells expressing FtsZ_DD_-GFP were compared with drFtsZ under both the growth conditions. The cells expressing FtsZ_DD_-GFP episomally showed cell division pattern similar to the cells expressing wild type protein episomally (Fig 4B). This result could be attributed to the dominant phenotype of chromosomally encoded drFtsZ over the FtsZ_DD_-GFP in wild type background. When exposed to γ-irradiation, both types of cells showed similar trend in cell division. Amazingly, the FtsZ_DD_-GFP expressing cells failed to form any new and active structures, instead they produced an erratic pattern of FtsZ_DD_ localization at 4h PIR (Fig 4B). Further, unlike the drFtsZ cells that got rescued from γ-irradiation effects in 4/5h of the PIR, the FtsZ_DD_ mutant failed to do so and continued to show aberrant pattern of localization even after several hours of the PIR period. These results suggested that the physiological behaviour of drFtsZ is similar to FtsZ_DD_ under normal growth conditions, possibly because of chromosomal copy of drFtsZ while grossly different under γ radiation stressed growth. These findings together suggested that the phosphorylation of drFtsZ at S235 and S335 positions by a γ radiation responsive STPK (RqkA) alters the typical characteristics of FtsZ and if that can arrest its dynamics in response to γ radiation exposure could be hypothesized.

### Mimicking phosphorylation affected *in vivo* dynamics of drFtsZ

A possible effect of phosphorylation on drFtsZ dynamics upon γ radiation exposure was studied by the fluorescence recovery after photobleaching (FRAP) approach. Wild type cells expressing the GFP fusion of both drFtsZ and FtsZ_DD_ mutant were also examined microscopically, and their foci were studied for FRAP. The results showed that the drFtsZ foci were able to recover under normal conditions (Fig 5A-B). In response to γ-irradiation, the bleached foci of drFtsZ could recover at 0h of PIR but not in 2h of PIR cells suggesting that the phosphorylation of FtsZ that would occur between 2 to 3h PIR under our experimental condition (Maurya et al., 2018) seems to have affected its polymerization/depolymerization characteristics *in vivo*. Notably, the bleached foci corresponding to FtsZ_DD_ protein did not recover under both normal and γ stressed growth conditions monitored under this study (Fig 5A-B). This might suggest that the phosphorylation of FtsZ regulates the dynamics needed for cells to undergo binary fission and thus could be the key regulator of cell cycle arrest upon γ radiation exposure at least in this bacterium.

**Fig 5:**
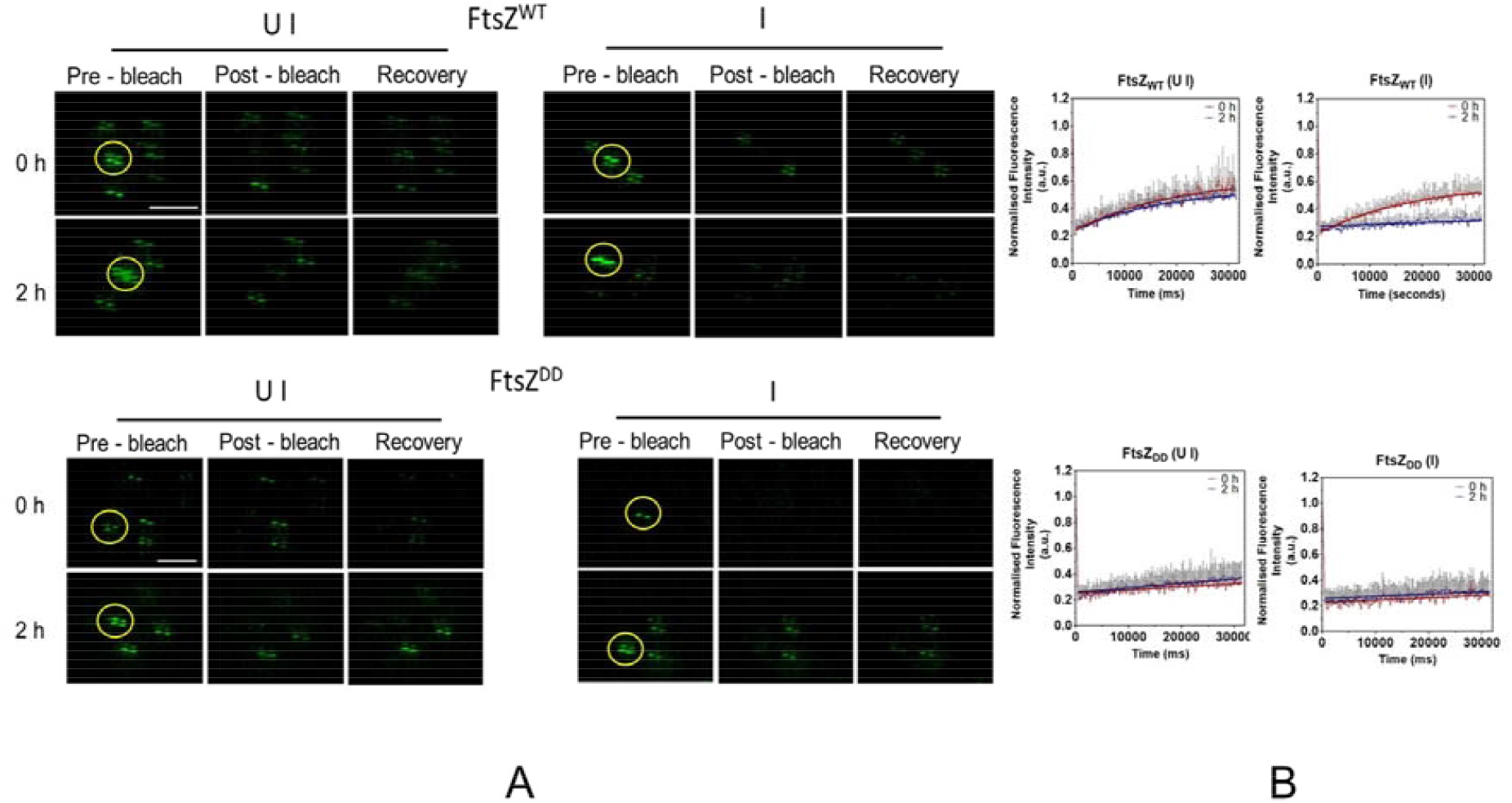
Mimicking phosphorylation mutation affect the recovery dynamics of drFtsZ. For FRAP microscopic analysis, the cells in exponential phase were exposed to 6 kGy dose of γ radiation whereas control cells were kept on ice. After γ-irradiation, both un-irradiated (UI) and irradiated (Irr) cells expressing pRGZGFP (FtsZ_WT_) and pRG_DD_GFP (FtsZ_DD_) mounted on 2X-TYG agarose pad before imaging. The representative images of pre-bleach, bleach and recovery phase are also shown **(A)**. FRAP-measurements were made at room temperature. The region of interest was exposed to laser bleach pulse of 35 to 45 ms followed by the acquisition of recovery for 30-40 s at low laser power. The fluorescence intensities in the bleached and other regions at each time point were extracted using the cellSens software and exported to Excel 2013. Recovery curves were plotted by performing a non-linear regression fit of the intensity of the bleached region over the time using GraphPad software **(B)**.

**Fig 6:**
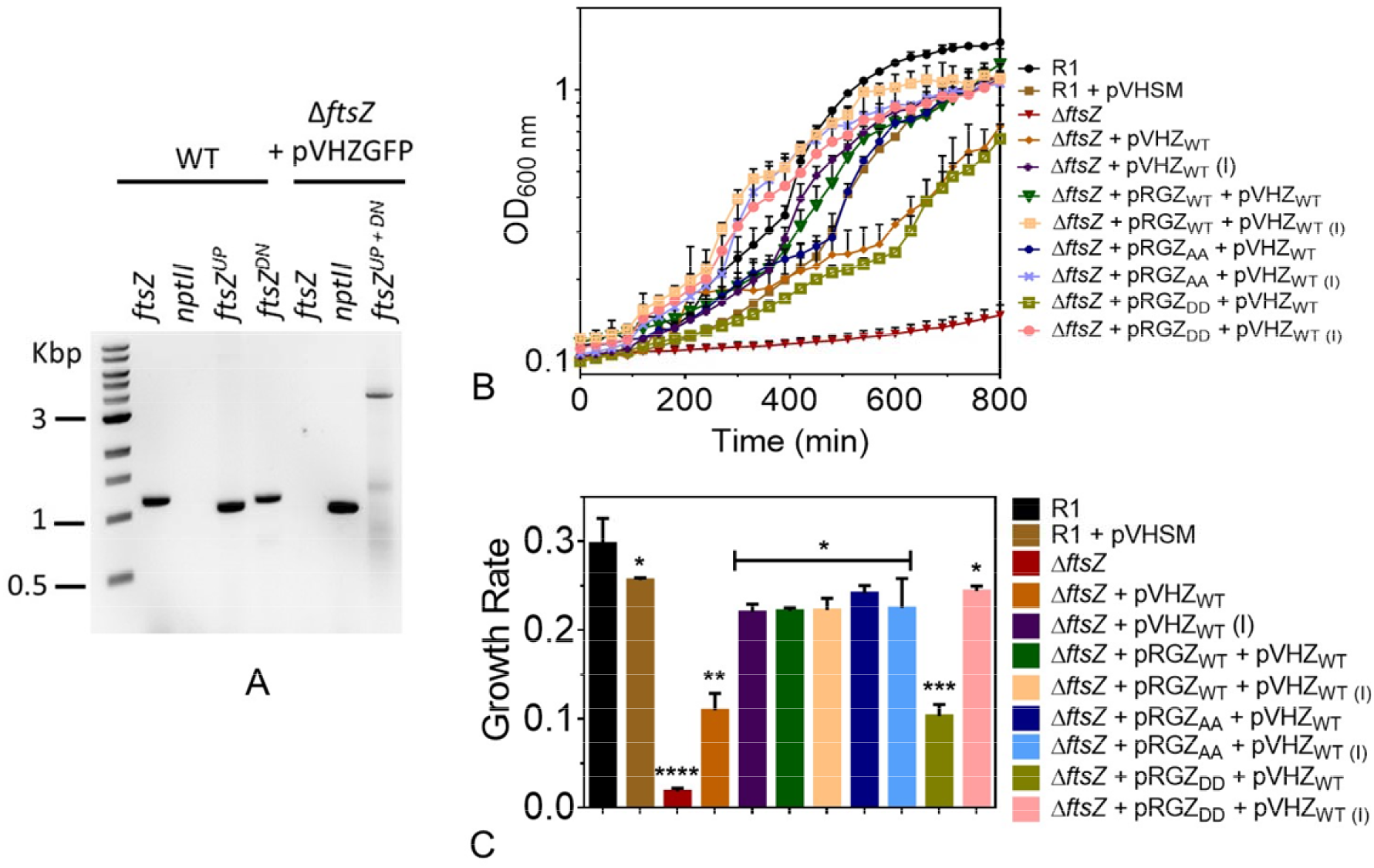
Phospho-mimetic mutant failed to support the cell-growth. The plasmid pNKZ was linearized and transformed into *D. radiodurans* cells harbouring pVHZGFP and transformants were scored in the presence of kanamycin and spectinomycin. The replacement of *ftsZ* with an antibiotic resistance marker gene (*nptII*) was confirmed by diagnostic PCR using primers as described in Table 2. The PCR products in the gel are labelled for their respective identity as following, *nptII* for antibiotic resistance gene, *ftsZ* for the FtsZ coding sequence, *ftsZ*^*UP*^ and *ftsZ*^*DN*^ for upstream and downstream sequences of *ftsZ*, respectively. The *ftsZ*^*UP + DN*^ indicates the product of *ftsZ*^*UP*^ *+ nptII + ftsZ*^*DN*^ from the conditional null mutant **(A)**. The wild-type and conditional null mutant derivatives were monitored for their growth kinetics using Synergy H1 multiplate reader. The wild-type strain (R1), R1 harbouring vector backbone of pVHSM (R1 + pVHSM), presumptive null mutant of *ftsZ* (Δ*ftsZ)* harbouring pVHZGFP (Δ*ftsZ* + pVHZ_WT_), conditional null mutant of *ftsZ* (Δ*ftsZ* + pVHZ^WT^) harboring pRGZGFP (pRGZ_WT_), pRGZ_AA_GFP (pRGZ_AA_) and pRGZ_DD_GFP (pRGZ_DD_) were monitored for their growth in the presence and absence of IPTG where I indicates the induced status of the pVHZGFP in the corresponding strain as shown in the graph **(B)**. The growth rate was calculated from the corresponding growth-curve values and plotted using GraphPad Prism software **(C)**.

### Phospho-mimetic mutant of drFtsZ failed to complement *ftsZ* loss in *D. radiodurans*

In order to understand the effect of S235 and S335 phosphorylation on growth in *D. radiodurans*, we needed the null mutant of *ftsZ*. Since the homogeneous deletion of *ftsZ* could not be possible, the *ftsZ* conditional-null mutant (Δ*ftsZ*) was created in the presence of episomal copy of drFtsZ. The homogenous replacement of target sequence with *nptII* cassette was confirmed by diagnostic PCR as shown in (Fig 5A). These cells grew normally and did not lose viability, and hereafter termed as Δ*ftsZ* conditional-null mutant (Δ*ftsZ*). The effect of phospho-mimetic replacement of S235 and S335 on functional complementation in Δ*ftsZ* was monitored under normal and γ radiation stressed conditions. For that, drFtsZ, FtsZ_AA_ and FtsZ_DD_ were expressed in Δ*ftsZ* cells on plasmids and the growth of these cells was monitored in the presence and absence of episomally expressed drFtsZ (Fig 5B). The Δ*ftsZ* cells expressing drFtsZ and FtsZ_AA_ on plasmid grew normal and nearly similar to wild-type cells irrespective of the episomal expression from drFtsZ suggesting that the basal expression of phospho-ablative mutant was enough for the normal growth of the Δ*ftsZ* cells. However, the Δ*ftsZ* cells expressing FtsZ_DD_ grew slower in the absence IPTG inducible expression of drFtsZ, which recovered when induced with IPTG. A minor growth in the Δ*ftsZ* cells expressing FtsZ_DD_ without IPTG may be accounted to the leaky expression of drFtsZ under IPTG inducible promoter. These results supported our hypothesis that mimicking phosphorylation at S235 and S335 in drFtsZ changing its polymerization/depolymerization characteristics, which is the key for bacterial cells to divide, and seems to be responsible for cell cycle arrest in response to γ radiation exposure in this bacterium.

## Discussion

FtsZ is a key protein in bacterial cell division and the GTP/Mg^2+^ supported polymerization/depolymerization dynamics is one of its intrinsic properties that regulate growth in bacteria. In response to DNA damage, the activity of FtsZ attenuated by SOS proteins like SulA in *E. coli*, (Mukherjee et al., 1998), YneA in *B. subtilis* (Kawai et al., 2003) and Rv2719c in *M. tuberculosis* (Chauhan et al., 2006). *D. radiodurans* is known for its extraordinary resistance to several DNA damaging agents including γ-radiation. The cells exposed to γ-radiation show the growth arrest for nearly 4 h, till presumably damaged DNA is repaired, despite no significant change in the levels of some cell division proteins. Unlike other bacteria, this bacterium does not deploy LexA/RecA type SOS mechanisms to control the cell cycle in response to DNA damage. However, it resists the lethal doses of γ radiation and other DNA damaging agents by both qualitative and quantitative changes in the genome functions. These findings collectively argued in the favour of some alternate mechanism/s of the DNA damage response and cell cycle regulation in this bacterium.

The post translation modifications (PTMs) is known to play the significant roles in the regulation of cellular dynamics of macromolecular interactions and stress responsive gene expression in higher organisms (Yakubu et al., 2018; Ivarsson & Jemth, 2019). Notably, Ser/Thr (S/T) phosphorylation of proteins is the key to DNA damage response and cell cycle regulation in eukaryotes. Recently, S/T phosphorylation has also been reported in bacterial proteins involved in the cell division, genome maintenance and pathogenesis (Rajpurohit et al., 2021). The S/T phosphorylation of DNA repair proteins like RecA has been demonstrated in *B. subtilis* grown under normal conditions (Bidnenko et al., 2013). The genome of D. radiodurans also encodes a few typical eukaryotic type STPKs and one such STPK (RqkA) has been characterised for its role in radioresistance. The *rqkA* mutant was found to be hypersensitive to γ radiation and showed near to complete arrest of DSB repair (Rajpurohit et al., 2010). RqkA could phosphorylate a number of deinococcal proteins including PprA, RecA, FtsZ and FtsA. The effect of S/T phosphorylation on the functions of DNA repair proteins (RecA and PprA) and radiation resistance has also been demonstrated in this bacterium (Rajpurohit et al. 2010; 2013). Although, the S/T phosphorylation of drFtsZ and drFtsA by RqkA, and their kinetic change during PIR has been reported (Maurya et al., 2018), the impact of such phosphorylation on the cell cycle regulation in response to DNA damage has not been shown yet. This study has brought forth some direct evidence to suggest that the phosphorylation of drFtsZ by RqkA has impacted its intrinsic properties *in vitro* and on the cellular dynamics under both normal and γ-stressed conditions *in vivo*. We demonstrated that the replacement of phosphorylated serines like S235 and S335 with aspartate reduces the GTPase activity by affecting GTP interaction with the protein. The polymerization ability of these mutants had been severely compromised *in vitro*, which appears to be due to the loss of the GTP binding ability of the protein. The effect of S235 and S335 phosphorylation on GTP binding pocket of drFtsZ was studied *in silico*. It was observed that the replacement of S235 or S335 or both with phospho-mimetic residue (aspartate) alters the position of the GTP binding pocket in drFtsZ (Fig S3A (i)). For instances, in drFtsZ, it was positioned in the region between 96 to 101 residues. Interestingly, in FtsZ_DD_, the GTP binding pocket got shifted to a region between 321 to 341 residues and become close to IDP territory (Fig S3A (ii)). In case of both single phospho-mimetic mutants; FtsZ_S235D_ and FtsZ_S335D_, the GTP binding pocket was also shifted to the region between 126 to 131 residues (Fig S3A (iii-iv)).

Although, the FtsZ phosphorylation has been reported in many bacteria including mycobacteria (Thakur & Chakraborti, 2006), streptococci (Giefing et al., 2010), staphylococci (Heumer et al., 2021) and *D. radiodurans* (Maurya et al., 2018), the phosphorylation of FtsZ affecting cell cycle in response to γ radiation is first time demonstrated in this bacterium. The localization pattern of phospho-mimetic mutant is found to be completely different from phospho-ablative mutant and wild-type in both *ex vivo* and *in vivo* conditions, suggesting that mimicking phosphorylation makes this protein compromising its native functions. Earlier it has been shown that phosphorylation of drFtsZ follows a kinetics during PIR with its highest levels during 2 h and 3 h PIR, which also coincides with the arrest of growth upon DNA damage. Therefore, a possibility of FtsZ phosphorylation arresting *in vivo* dynamics and thereby growth arrest can be hypothesized. The FRAP analysis of wild-type and phospho-mimetic mutant clearly supported our assumption that the dynamics of phosphorylated FtsZ gets arrested upon γ exposure. To the best of our knowledge, the involvement of S/T phosphorylation on FtsZ dynamics has not been shown before in any bacteria and therefore, it offers a unique example of LexA/RecA independent cell cycle arrest through S/T phosphorylation in this bacterium.

In conclusion, this study has provided some evidence to suggest that the DNA damage response and cell cycle in response to γ radiation is regulated by eukaryotic type STPK (RqkA) in *D. radiodurans* that apparently lacks the canonical SOS response as known in other bacteria. Independently, it has been shown that S/T phosphorylation of DNA repair proteins stimulates their functions and is needed for their support to radioresistance. Here, we have provided evidence that the mimicking the phosphorylation of S235 and S335 of FtsZ causes the loss of its polymerization ability *in vitro* and the arrest of cellular dynamics *in vivo*. The arrest of drFtsZ dynamics has also been observed in *D. radiodurans* in response to γ radiation but this showed reversal at a later phase of PIR. However, the phospho-mimetic mutant did not resume dynamics during the PIR period monitored under this study. These findings allow us to conclude that S235 and S335 phosphorylation in drFtsZ by RqkA in response to γ radiation seems to be the key regulator of FtsZ functions and cell cycle regulation in this bacterium.

## Acknowledgements

Authors would like to thank Mrs. Rimanshee Arya for her help with the Malvern Zetasizer and JASCO Spectrophometer and Mr Shishu Kant Suman in optimization for using fluorescent labelled nucleotides. Reema Chaudhary is grateful to Department of Atomic Energy, India for her fellowship.

## Declaration

This work has also been a part of doctoral thesis of Dr. Reema Chaudhary.

---

**Fig S1:**
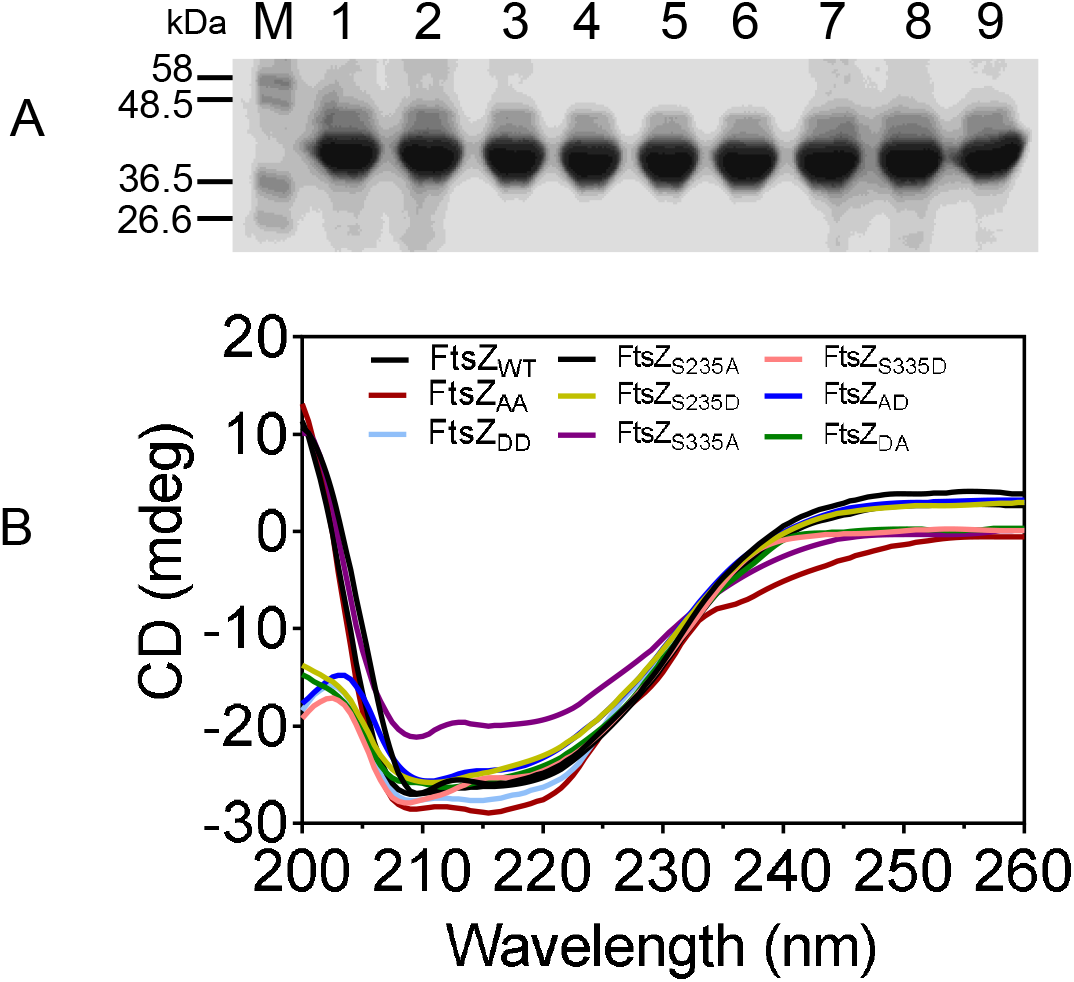
Purification of phospho-mutants of FtsZ and their CD spectra. The phospho-mutants of FtsZ (1: FtsZ_WT_, 2:FtsZ_AA,_ 3:FtsZ_DD,_ 4:FtsZ_S235A,_ 5:FtsZ_S335A,_ 6:FtsZ_S235D,_ 7:FtsZ_S335D,_ 8:FtsZ_AD,_ 9:FtsZ_DA)_ were purified under native conditions and checked on SDS-PAGE **(A)**. CD Spectra of FtsZ and its phospho-mutants was monitored and the proteins were diluted in buffer containing 20 mM Tris (pH 7.6) and 150 mM NaCl and spectra was recorded using CD spectrophotometer (Biologic spectrometer MOS-500). The CD spectra of the wild type and phospho-mutant derivatives of FtsZ have shown the typical CD spectra suggesting the formation of correct secondary structures.

**Fig S2:**
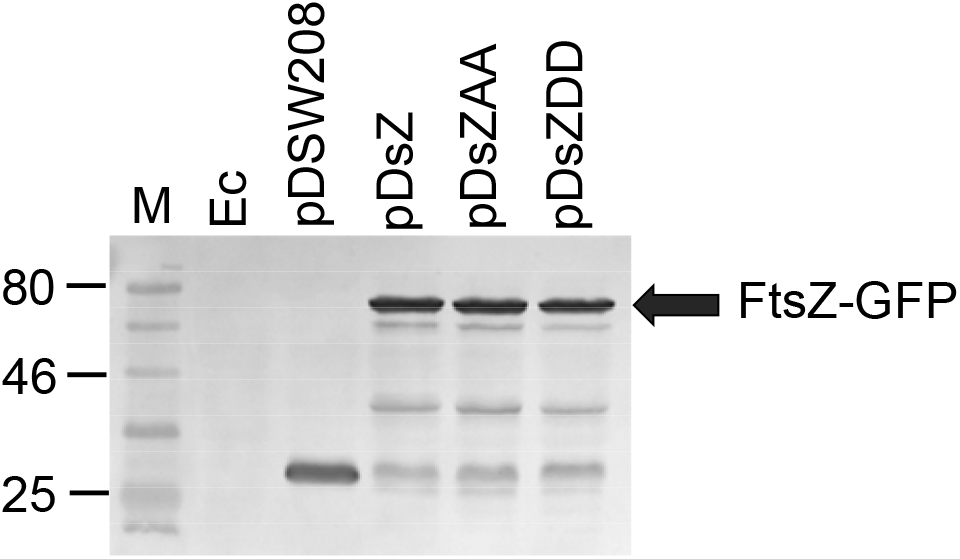
The total proteins of *E. coli* (Ec) harboring pDSW208, pDsZ, pDsZAA and pDsZDD were separated on SDS-PAGE and immunoblotted with anti-GFP antibodies.

**Fig S3:**
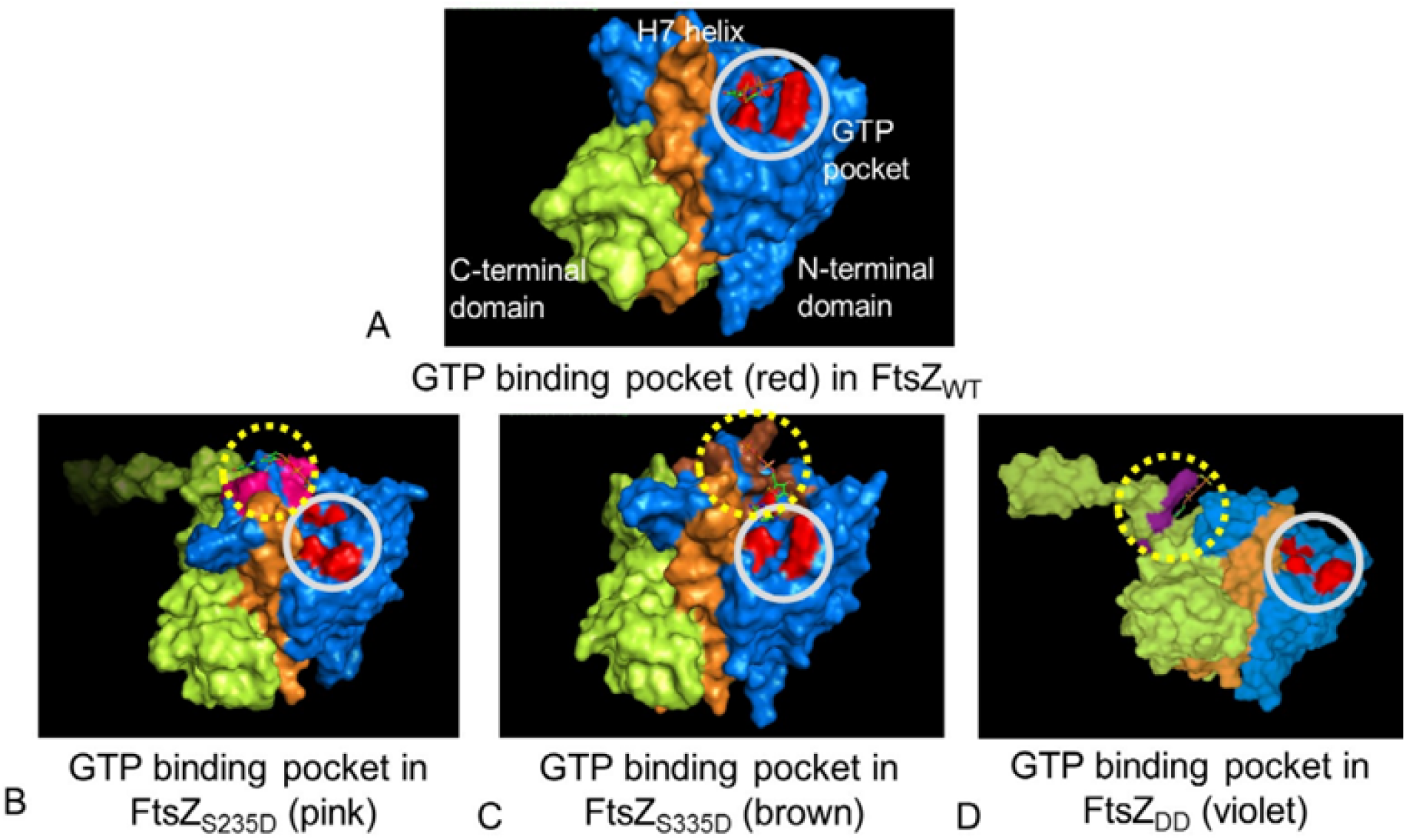
Mapping of GTP-binding pocket in the wild type and phospho-mimetic mutants of FtsZ using an *in-silico* approach. The structures of FtsZ_WT_, FtsZ_S235D_ FtsZ_S335D_ and FtsZ_DD_ were modeled using an I-TASSER online tool and used for docking with GTP using an Autodock tool and the program was run for predicting the position of its binding in the protein. The structure of the proteins is color-coded as blue for its N-terminal domain, orange for H7-helix and bright green for the C-terminal domain. The GTP binding pocket in FtsZ_WT_ is marked as red (A) which shifts to other positions in mutants, shown by pink, brown and violet color in FtsZ_S235D_ (B) FtsZ_S335D_ (C) and FtsZ_DD_ (D). Further, for more clarity, the nucleotide-binding pocket is highlighted with a gray-colored solid circle for depicting its wild-type position while the change in its position is shown with a yellow-colored dotted circle in phospho-mimetic proteins.

